# Heterogeneous Tissue Modulus Improved Prediction of Mechanical Behavior in Osteoporotic Vertebral Cancellous Bone

**DOI:** 10.1101/2021.11.30.470675

**Authors:** Jason M. Cox, Joshua D. Smith, Marjolein C. H. van der Meulen, Jacqueline H. Cole

**Author notes:** Corresponding Author, J. H. Cole. M. C. H. van der Meulen.

## Abstract

The structural integrity of cancellous bone, which is essential to its skeletal load-bearing capacity, is governed chiefly by apparent density, trabecular architecture, and tissue material properties. Metabolic bone disorders such as osteoporosis can affect each of these factors, resulting in compromised load-bearing function and fracture. While the impact of apparent density and architecture on bone structural behavior is well-documented, much less is known about the influence of tissue material properties, particularly in osteoporotic bone. In this work, we isolated the influence of tissue modulus on normal and osteoporotic cancellous bone structural integrity, indicated by the apparent elastic modulus under uniaxial compression and patterns of internal tissue strain. Finite element (FE) models derived from 3D micro-computed tomography images were compared to physical testing data of the same samples. Three sets of FE models with increasing material detail were studied: 1) universal tissue elastic modulus (20 GPa), 2) specimen-specific average tissue modulus, and 3) heterogeneous tissue modulus. Applying a universal modulus resulted in overestimation of osteoporotic bone apparent modulus; applying specimen-specific material properties, either as a single average tissue modulus or heterogeneous distribution of tissue moduli, prevented significant apparent modulus overestimation. The greatest improvement in apparent modulus prediction resulted from incorporating a specimen-specific average tissue modulus, though using a specimen-specific heterogeneous tissue modulus provided the most reliable prediction of apparent modulus overall. In addition, median element strain in heterogeneous models also trended lower than in homogeneous models. This finding suggests that heterogeneous material properties may play a role in protective strain-concentrating mechanisms observed in cancellous bone. We conclude that future work exploring trabecular bone mechanics through finite element analysis should incorporate specimen-specific average tissue modulus at a minimum, but heterogeneous tissue modulus is recommended to maximize the functional similarity of bone *in* silico with bone *in vivo*.

## 1. Introduction

The structural integrity of cancellous bone, which is essential to skeletal load-bearing capacity, is governed chiefly by apparent density, trabecular architecture, and tissue material properties (Arlot et al., 2008; Beaupre and Hayes, 1985; Carter and Hayes, 1976; Cory et al., 2010; Goldstein, 1987; Martin et al., 1998; Raux et al., 1975; Townsend et al., 1975). Although the impact of apparent density and architecture on bone structural behavior has been well-documented, less is known about the influence of tissue material properties. Bone tissue is innately heterogeneous, exhibiting substantial spatial variation in tissue mineral content (Bourne et al., 2002; Morgan et al., 2002; Paschalis et al., 1996, 1997a). Metabolic conditions such as osteoporosis affect bone tissue properties, as generally evidenced by altered mineralization and changes in chemical composition (Gadeleta et al., 2000; Grynpas, 1993; Miller et al., 2000; Paschalis et al., 1997b). Changes in tissue properties may compromise the structural performance of bone, although the specific contributions of tissue mineralization and composition to bone mechanical properties have not been fully elucidated, particularly for cancellous bone.

Tissue properties are difficult to characterize experimentally and are primarily examined in small, excised bone volumes that likely do not fully capture the *in vivo* spatial variability. Computer models that simulate a virtual biopsy of cancellous bone can be used to investigate the mechanical behavior of a much larger region of bone and provide the opportunity to analyze that behavior noninvasively over changes in structural, material, and external loading parameters. Early finite element (FE) models treated cancellous bone as an irregular lattice-type structure and assessed the effect of idealized changes in architecture but not tissue properties (Beaupre and Hayes, 1985; Guo et al., 1994; Jensen et al., 1990; Silva and Gibson, 1997; Yeh and Keaveny, 1999). Architecture-based models, developed by converting voxels from a high-resolution micro-computed tomography (micro-CT) scan into hexahedral finite elements, more closely mimic the bone structure with fewer geometric assumptions than in the lattice models (Feldkamp et al., 1989). In the past, these voxel-based models were primarily used to examine the role of architecture in cancellous bone mechanical behavior through finite element analysis (FEA). Most models assumed isotropic and homogeneous material properties (Bevill et al., 2009; Chevalier et al., 2007; Hollister et al., 1994; Hou et al., 1998; Kabel et al., 1999a, 1999b; Ladd et al., 1998; Liu et al., 2006; Niebur et al., 2001, 2000; Ulrich et al., 1997; Unnikrishnan et al., 2015; van Rietbergen et al., 1998, 1997, 1995). More recently, voxel-based modeling studies have investigated the interplay of cancellous bone mechanics and physiological processes such as microdamage progression (Goff et al., 2015; Liu et al., 2009; Torres et al., 2019), homeostatic bone remodeling (Hambli et al., 2011; McNamara and Prendergast, 2007; Pauchard et al., 2008), and bone remodeling following implantation of an orthopaedic device (Li et al., 2019; Schulte et al., 2013). Although these processes occur on a tissue-level scale at which variable material properties may play a significant role, the assumption of material homogeneity is still common. Another approach to architecture-based FE modeling involves the use of beam elements to model whole trabeculae, dramatically reducing model degrees of freedom and thus computational demands (van Lenthe et al., 2006; Wang et al., 2019). While this strategy is advantageous for analysis throughput and has produced accurate predictions of whole-bone level mechanical behavior, this approach prohibits the inclusion of physiological material heterogeneity and the investigation of tissue behavior at the sub-trabecular level.

Material heterogeneity in bone tissue manifests as variable tissue mineral density (*ρ*), which can be converted to an elastic modulus (*E*) through empirically derived relationships and assigned to elements in the models. *E*-*ρ* relationships vary across species and bone sites (Beaupre and Hayes, 1985; Goulet et al., 1994; Helgason et al., 2008; Hodgskinson and Currey, 1992; Oftadeh et al., 2015; Rice et al., 1988) but generally take a power law form (Currey, 1986). Early efforts to incorporate tissue heterogeneity in FE models imposed assumed mineral distributions approximating natural tissue heterogeneity. These efforts revealed that the apparent elastic modulus of cancellous bone with tissue heterogeneity could be higher or lower than that of the same structure with homogeneous tissue, depending on the form of the power law used (van der Linden et al., 2001). Furthermore, increasing tissue modulus variability correlated with decreased apparent elastic modulus and an increased percentage of bone elements exhibiting elevated strain (Jaasma et al., 2002). As a more modern alternative to assuming the tissue mineral distribution, calibrated micro-CT scans provide an accurate representation of tissue mineral distribution (Homminga et al., 2001) that can be readily converted into a tissue modulus distribution through empirically derived *E*-*ρ* relationships. Several previous studies employed this method to investigate the effect of tissue mineral heterogeneity on cancellous bone behavior; however, a conclusive effect in human tissues was not demonstrated, because these studies used data from a different species (Bourne and van der Meulen, 2004; Nagaraja et al., 2007), a small sample size (Hammond et al., 2018; Nagaraja et al., 2007; van Ruijven et al., 2007), an entirely male cohort (Renders et al., 2011, 2008), or homogeneous models that did not match the average density of the heterogeneous test case, confounding comparisons between the two (Knowles et al., 2019; Nagaraja et al., 2007).

The question remains as to whether physiological heterogeneity in FE models, and by extension *in vivo*, has a substantial influence on human cancellous bone mechanical behavior. In addition, how the influence of heterogeneity depends on sex or the presence of metabolic bone disorders such as osteoporosis is not fully understood. Given that osteoporosis alters bone mineralization and osteoporosis prevalence differs between men and women, these parameters need to be examined to assess the robustness of FE models for predicting cancellous bone behavior. The objectives of this study were 1) to simulate the structural behavior of cancellous bone in the human spine using specimen-specific architecture-and material-based FE models; 2) to assess the effects of varied material models on the mechanical behavior of the bone structure; 3) to compare model-derived parameters to experimentally measured mechanical properties of the same cancellous specimens; and 4) to determine the effects of sex, bone site, and clinical bone diagnosis on the mechanical behavior of the cancellous specimens across the various material models. Our approach was to model the architecture and material variations of human vertebral cancellous bone based on micro-CT scans, perform finite element analyses on the models, and compare the model predictions for different material distributions to experimentally measured mechanical properties of the same specimens.

## 2. Materials and Methods

### 2.1 Subjects

Cadaver spine segments were obtained from 19 Caucasian donors (10 female, 9 male) ages 58 to 92 years. For 18 donors, the spine segment extended from the 11^th^ thoracic vertebra (T11) through the 4th lumbar vertebra (L4), and for one donor, the segment extended from T11 through the 2nd lumbar vertebra (L2). The spines had no evidence of vertebral fracture in a clinical bone density scan or of bone abnormalities during dissection, such as bone metastasis.

### 2.2 Clinical Bone Density Scan

Clinical bone mineral density status was assessed for all spines with dual-energy X-ray absorptiometry (DXA). Spine segments were secured in a curved Plexiglas® fixture, immersed in a saline bath within a Plexiglas® box, and scanned with a clinical fan-beam densitometer in lumbar spine array mode (Delphi QDR 4500A or QDR 4500W, Hologic Inc., Bedford, MA). T-score was computed for the L1-L4 subregion of each spine using the Hologic analysis software, and subjects were classified by T-score as Non-osteoporotic (T-score > -2.5) or Osteoporotic (T-score ≤ -2.5), according to the World Health Organization guidelines (Kanis et al., 1994).

### 2.3 Micro-Computed Tomography Scan

To prepare small cancellous bone specimens for micro-CT, the T12 and L2 vertebrae were excised from each spine segment, and a full-depth cylindrical core (nominal diameter = 8.25 mm) was obtained from the center of each vertebra. The vertebra was secured in a drill press, and the cut was made along the superior-inferior axis at low speed under a continuous saline stream using a metal bond diamond core drill (Starlight Industries, Rosemont, PA). Cored specimens were wrapped in saline-soaked gauze and stored frozen at -20°C until scanning.

To obtain a voxel-based characterization of cancellous bone structure and mineralization, cored vertebral specimens were scanned using quantitative micro-CT (MS-8, GE Healthcare, London, Ontario, Canada) with 70 kVp X-ray tube peak potential, 90 μA X-ray intensity, and 3 s integration time. The specimens were scanned in saline treated with a protease inhibitor cocktail (P8340, Sigma-Aldrich Inc., St. Louis, MO) to minimize protein degradation in the bone tissue and then refrozen at -20°C until mechanical testing. The scans were reconstructed at an isotropic voxel size of 17.0 μm, and mineral density was calibrated using a phantom consisting of air, water, and a bone mineral standard (SB3, Gammex RMI, Middleton, WI) with a known density of 1.15 g/cc. For analysis, the scans were reoriented to align the superior-inferior axis of the core with the vertical axis, and the ends of the image were cropped flat to a length of 17 mm (MicroView ABA 2.1.1., GE Healthcare, London, Ontario, Canada). Past work has suggested that the use of a global segmentation threshold for a cohort with a broad range of bone volume fraction or varying bone mineral density can result in an inaccurate topology (Bouxsein et al., 2010). To account for the inclusion of both osteoporotic and non-osteoporotic subjects in our cohort, specimen-specific thresholds were determined using the Otsu method (N. Otsu, 1979), and average tissue mineral density (TMD) was assessed for each core.

### 2.4 Mechanical Testing

Following micro-CT scans, the cancellous bone cores were tested destructively in uniaxial compression, and apparent-level properties were assessed. After the cortical end plates were removed, the bone core ends were glued inside snug-fitting brass caps using a previously documented protocol designed to minimize end artifacts (Keaveny et al., 1994). Approximately one-fourth of the overall core length was capped at each end. The diameter of each cancellous core was measured six times, and the exposed length was measured four times, using metric dial calipers, and the mean values were recorded. After the cyanoacrylate glue was cured for 24 hours at 4°C, the capped cores were preconditioned for 5 cycles between 0 and 0.10% compressive strain and loaded monotonically to failure at a rate of 0.50% compressive strain per second (Mini-Bionix 858, MTS Systems Corporation, Eden Prairie, MN). Tests were conducted in displacement control at room temperature, and load and displacement data were sampled at 20 Hz. Displacement was measured with a 25-mm gage length axial extensometer (634.11F-24, MTS Systems Corporation, Eden Prairie, MN) attached to both brass caps. To compute apparent mechanical properties, load and displacement data were converted to apparent stress and strain using standard mechanics formulae (Hibbeler, 2016). The experimentally measured apparent modulus (E_meas_) was defined as the slope of a least-squares linear fit to the elastic region of the stress-strain data over 0.02-0.24% strain (Kopperdahl and Keaveny, 1998).

### 2.5 Finite Element Modeling

The micro-CT scans were used to build specimen-specific finite element (FE) models for the bone cores (Hollister et al., 1994; van Rietbergen et al., 1995). Using the thresholded micro-CT scans, bone voxels were converted into 8-noded linear hexahedral elements, creating a FE mesh measuring 8.25 mm in diameter and 17 mm in length (**Figure 1**). Elements unconnected to the primary structure were removed using the connectivity criterion of a 6-connected neighborhood. Unless a face was shared with a portion of the primary bone structure, an element was considered unconnected and removed from the model. For all models, the number of unconnected elements removed was between 0.6% and 7.6% of the total number of elements. On average, the final FE models had 24,000,000 elements and 33,000,000 nodes.

**Figure 1:**
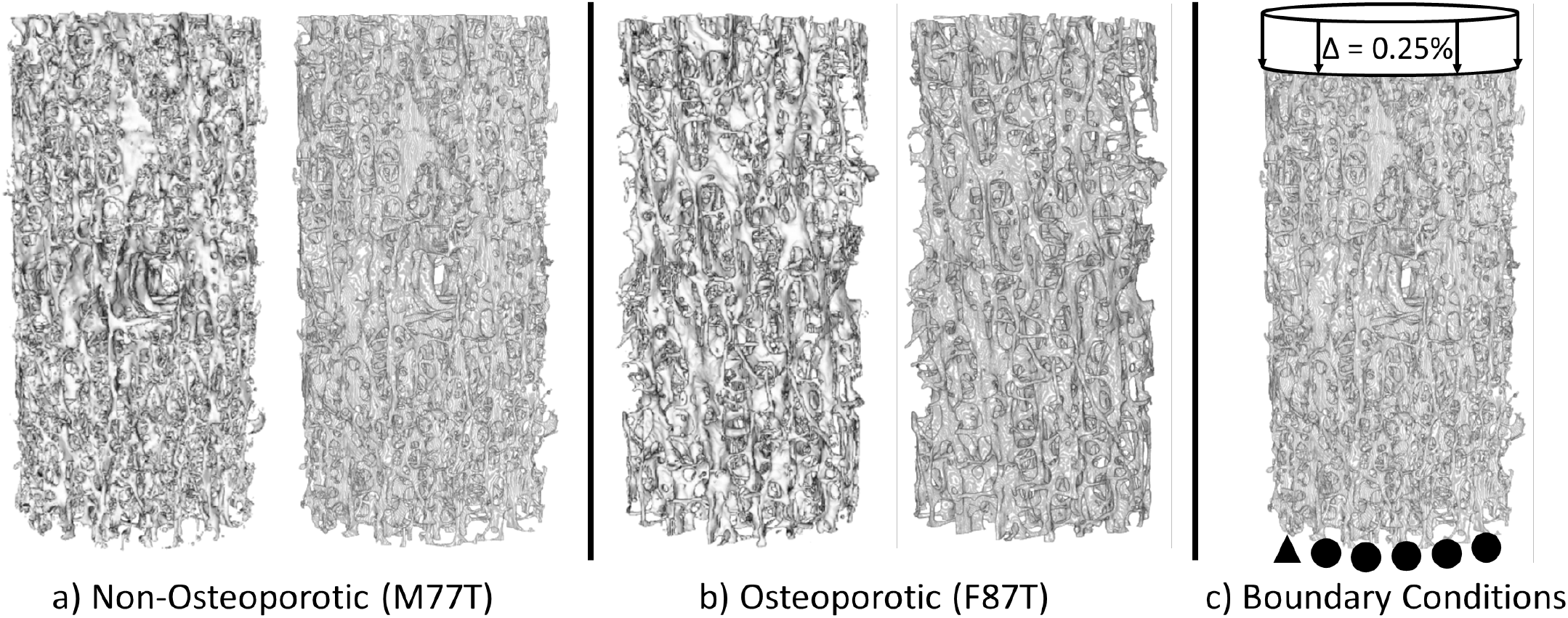
Examples of reconstructed volumes (left) and finite element meshes (right) for (a) non-osteoporotic and (b) osteoporotic subjects. Sex (M=male, F=female), age in years, and vertebral site (T=thoracic) are denoted. (c) Boundary conditions for finite element models. A compressive strain of 0.25% was applied to nodes on the top surface of the mesh. Nodes on the bottom surface were constrained to in-plane motion, except two adjacent nodes on the edge that were completely constrained to prevent rigid-body motion.

Three sets of FE models were created for each bone core using different material properties. The model sets were assigned isotropic material properties with a constant Poisson’s ratio of 0.3 and one of the following three tissue modulus distributions:

1. Universal tissue modulus (E_U_): A single homogeneous modulus of 20 GPa, chosen based on measurements from nanoindentation studies (Rho et al., 1999; Wang et al., 2006), was assigned to all elements for all bone core models.
2. Specimen-specific average tissue modulus (E_SSavg_): A unique homogeneous modulus was applied to all elements of each bone core model, computed using the mean voxel tissue density from the micro-CT scan of that bone core and a linear modulus-density relationship (Bourne and van der Meulen, 2004), as follows:

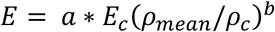

where *Ec* is the compressive tissue modulus for fully mineralized cortical bone (defined as 20 GPa), *ρ*_*mean*_ is the mean density over all mineralized voxels of the bone specimen, *ρ*_*c*_ is the density of the SB3 cortical bone mineral standard, and *a* and *b* are, respectively, the linear scaling factor and the exponent defining the nature of the *E-ρ* relationship (i.e., *b* = 1 for a linear relationship). Using a linear relationship between voxel tissue density and CT attenuation values and a linear scaling factor of 1.4 (Bourne and van der Meulen, 2004), the *ρ*_*mean*_/*ρ*_*c*_ ratio was computed using the ratio of the mean CT attenuation value to the SB3 mineral standard CT value.
3. Specimen-specific heterogeneous tissue modulus (E_SShet_): A unique heterogeneous modulus was computed for each element within each bone core model, using the same linear *E-ρ* formula, except *ρ*_*mean*_ was replaced by the individual voxel tissue density *ρ*_*voxel*_, and *E* was computed individually for each element using the CT value for the associated voxel. Element moduli were grouped into ∼1,000 equally spaced bins to simplify material mapping while maintaining ample variability.

To account for scanning artifacts, which appear as abnormally high-density regions in the bone, ranges of measured tissue density for each model were computed and normalized to the value of *ρ*_*c*_. These ranges were used to determine a global upper threshold of relative tissue density, and voxels with tissue densities above this threshold were assigned the mean value of tissue densities computed from the remaining voxels. Only three models were affected by this procedure, with less than 0.002% of elements affected for each. Applying this correction prevented the rare scanning artifacts from artificially decreasing the degree of heterogeneity in the models by increasing the width of element modulus bins.

Boundary conditions were applied to the models to replicate displacement-controlled uniaxial compressive loading in the linear elastic range (**Figure 1c**). Pre-yield compression was simulated by displacing the nodes on the top surface of the cancellous core mesh to a strain of -0.25%. The nodes on the bottom surface of the mesh were treated as rollers constrained to in-plane motion only, and two adjacent nodes at one edge of the bottom surface were rigidly fixed to prevent rigid-body motion. The FE models were solved using a linear elastic static analysis (ABAQUS 2018, Dassault Systèmes, Vélizy-Villacoublay, France), and the model-predicted apparent modulus and distribution of minimum principal strains throughout the bone structure were obtained. The apparent modulus was calculated by summing the nodal reaction forces on the bottom surface of the core mesh and dividing by the cross-sectional area of a circle enclosing that surface. Additionally, the percent error in model-predicted apparent modulus relative to experimentally measured apparent modulus was computed. Minimum principal strain (*ϵ*_*1*_) was analyzed at the element centroids throughout the model, and strain values were categorized as low (*ϵ*_*1*_ < 500 microstrain), middle (500 microstrain ≤ *ϵ*_*1*_ < 1500 microstrain), or high (*ϵ*_*1*_ ≥ 1500 microstrain), and the proportions of elements belonging to each category (%Low, %Mid, %High) were determined. Because the distributions of *ϵ*_*1*_ had a distinctly negative skew, with most of the values concentrated near zero, the median was used as the measure of centrality in lieu of the mean.

### 2.6 Statistical Analyses

All statistical models were constructed with a significance level of 0.05 (SAS University Edition v. 9.4, SAS Institute Inc., Cary, NC). Statistical models were used for the following analyses:

#### Apparent Modulus

1) The predictive effects of bone volume fraction (BV/TV), mean TMD, and TMD coefficient of variation (CV) on measured apparent modulus were investigated with a mixed linear model. A preliminary correlation analysis with BV/TV, mean TMD, and TMD CV as fixed effects revealed that mean TMD and TMD CV were highly correlated (r = -0.93). Therefore, a Gram-Schmidt orthonormal factorization of TMD CV was performed to isolate the orthogonal component relative to mean TMD, u_TMDCV_. Then a mixed linear model was constructed with BV/TV, mean TMD, and u_TMDCV_. 2) Apparent modulus was compared between groups stratified by estimation method (i.e., mechanical testing and three sets of FE models), sex, vertebral site, and clinical bone diagnosis using a mixed linear model. A preliminary model with full effects revealed that interactions between sex, vertebral site, and clinical bone diagnosis were not significant, and these interaction effects were not included in the final model to limit the number of comparisons. Post hoc pairwise comparisons between modulus estimation methods were performed separately by sex, site, and clinical bone diagnosis.

#### Apparent Modulus Prediction

1) The effects of model material distribution, clinical bone diagnosis, and their interaction on the absolute value of modulus prediction error (i.e., error magnitude) were investigated with a mixed linear model. Fixed effects in this analysis did not include sex and vertebral site, because initial results from the preceding analyses indicated that apparent modulus did not differ along these strata. 2) The mean signed modulus prediction error (i.e., error bias) was compared to a hypothetical mean of 0 using one-sample t-tests for each combination of clinical bone diagnosis and model material distribution. 3) Signed modulus prediction error for osteoporotic vs. non-osteoporotic models was compared for each material distribution using two-sample t-tests. 4) The proportion of variance in measured modulus explained by FE-predicted modulus (r^2^) was determined through simple linear regression for each modeled material distribution. Then, clinical bone diagnosis was added as a second explanatory variable, and multiple linear regression was performed for each modeled material distribution to examine the resulting improvement in r^2^.

#### Minimum Principal Strain

Element strain patterns were compared between groups stratified by FE model material distribution, sex, vertebral site, and clinical bone diagnosis. Separate mixed linear models were constructed for median, standard deviation (SD), and %Low, %Mid, and %High *ϵ*_*1*_ as the outcome variables. By the same rationale as apparent modulus analysis #2, the effects of sex and clinical bone diagnosis were not included, as a preliminary model with full effects indicated they were not significant. For each *ϵ*_*1*_ statistic, post hoc comparisons by FE model material distribution were also performed for subsets of the cohort separated by site.

To correct for each donor spine being represented twice in the dataset (for T12 and L2 vertebrae), donor ID was incorporated in all linear models as a random intercept effect. For each mixed linear model, the Benjamini-Hochberg correction was applied to prevent multiple tests for effects from inflating the false discovery rate (Benjamini and Hochberg, 1995). Likewise, for apparent modulus prediction analyses #2 and #3 and each group of post hoc comparisons, the Tukey-Kramer correction was applied to prevent multiple comparisons from inflating the false discovery rate.

## 3. Results

CT-based finite element models were developed for thoracic and lumbar vertebral cancellous bone cores of 19 subjects (10 female, 9 male), ages 58-92, with a broad range of DXA T-score (**Table 1**). Across samples, BV/TV ranged from 5.8% to 14.1% with a CV of 21%. Mean tissue mineral density from micro-CT varied slightly less, ranging from 0.56 to 0.85 g/cc, with a CV of 13%. TMD variability throughout the reconstructed bone volumes, assessed by computing the CV of normalized CT attenuation across all bone voxels, ranged from 18.5% to 37.1%. Neither BV/TV, mean TMD, nor TMD CV differed significantly by subject sex, bone site, or clinical bone diagnosis. For models with specimen-specific heterogeneous material properties, the distribution of bone tissue modulus was identical to that of tissue mineral density, given their linear relationship, and was similar in shape for all subjects (**Figure 2**). Overall, the heterogeneous tissue modulus averaged 14.5-22.2 GPa over all elements for a given model (**Table 1**), resulting in a global mean of 18.5 GPa over all models.

**Figure 2:**
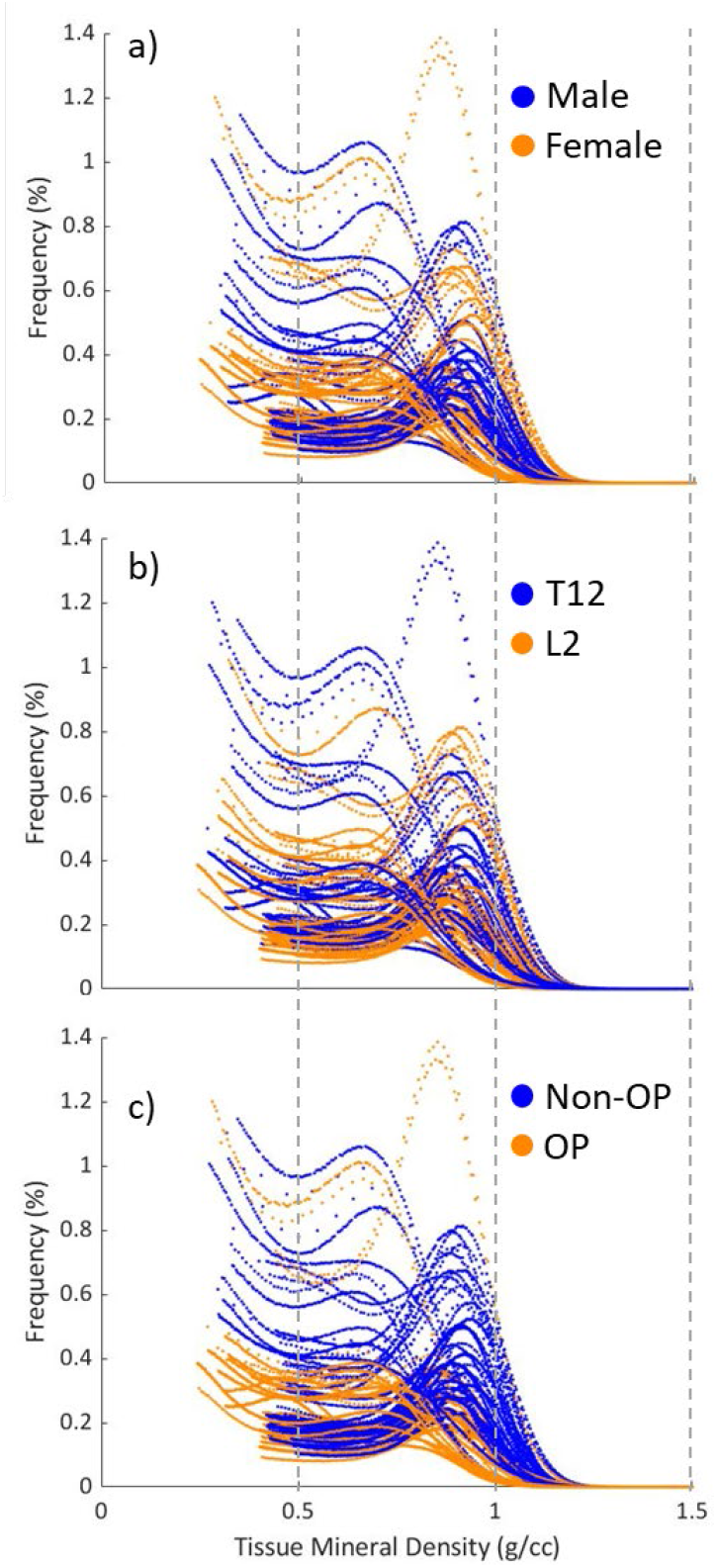
Bone tissue mineral density distribution assessed using CT attenuation in T12 and L2 bone cores for all 19 subjects, separated by (a) sex, (b) vertebral site and (c) clinical bone diagnosis (OP = osteoporotic). Shown are the percentages of voxels for each CT attenuation value, normalized by the CT value for the cortical bone phantom and scaled by its density (1.15 g/cc). Only voxel densities above the bone threshold are shown for each subject.

**Table 1:**
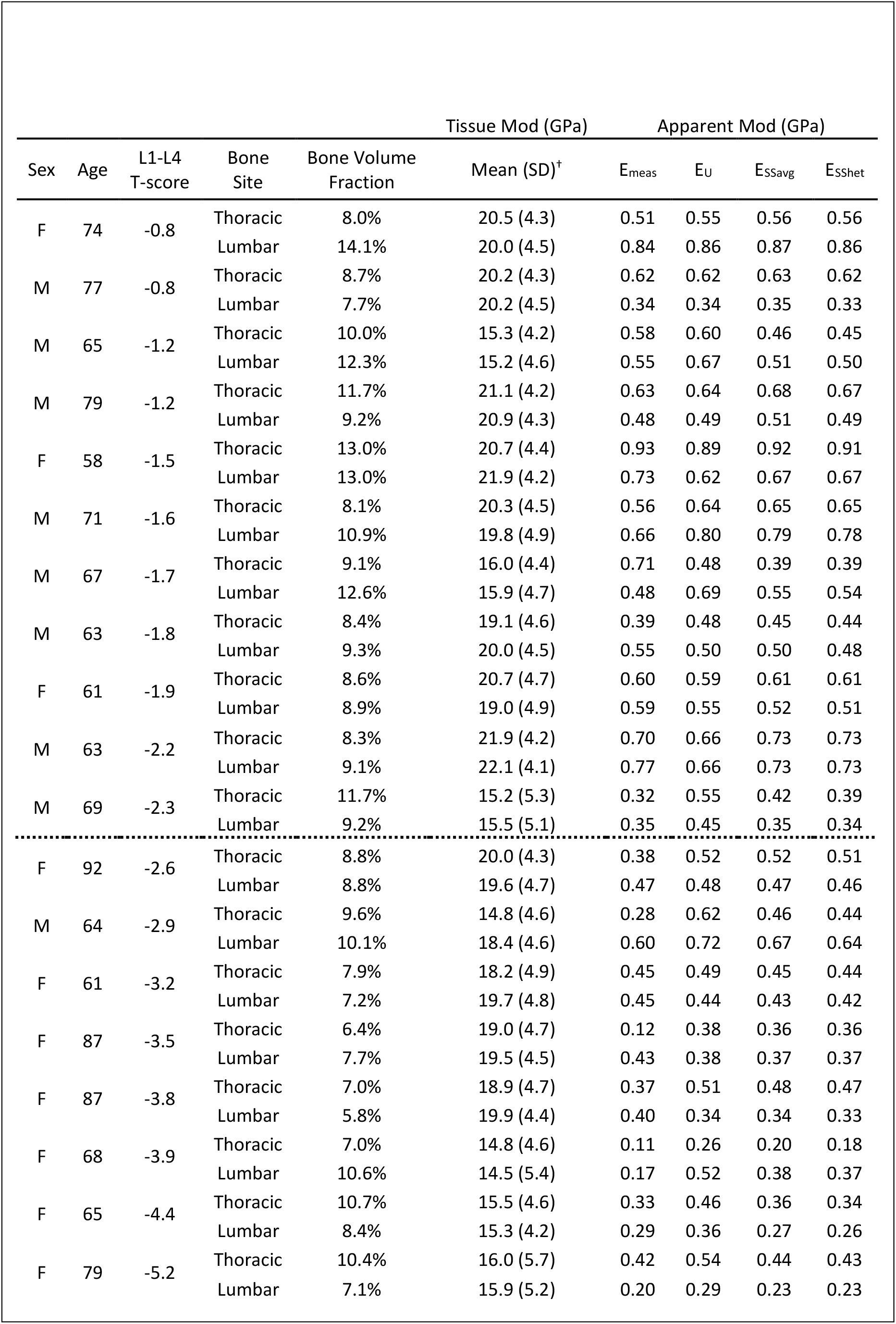
Mean tissue modulus (mod) for specimen-specific (SS) average (avg) and heterogeneous (het) models and mean apparent modulus measured experimentally (E_meas_) and for all models. Universal tissue modulus (EU) models used a homogeneous value of 20 GPa. Dotted line separates non-osteoporotic subjects (above) from osteoporotic subjects (below).

### Apparent Modulus

Experimentally measured apparent modulus ranged 0.11-0.93 GPa and was positively associated with BV/TV and mean TMD (p < 0.0001 for both), whereas for a given value of mean TMD, TMD CV had only a marginal negative association with measured modulus (p = 0.061), as indicated by u_TMDCV_. Under uniform compressive strain, FE-predicted apparent modulus was similar across the three model sets, ranging 0.26-0.89 GPa for E_U_ models, 0.20-0.92 GPa for E_SSavg_ models, and 0.18-0.91 GPa for E_SShet_ models (**Table 1, Figure 3a**). Mean FE-predicted apparent modulus across all samples was not significantly different from the mean measured apparent modulus for any of the three assigned tissue modulus distributions (p = 0.11). Regardless of the estimation method, apparent modulus differed by clinical bone diagnosis, with 36% lower apparent modulus for osteoporotic compared to non-osteoporotic bone (0.38 vs. 0.60 GPa, p = 0.007, **Figure 3a**). Within the osteoporotic subset, E_U_ models had a significantly higher mean apparent modulus compared to the mean experimentally measured modulus (0.46 vs. 0.31 GPa, p = 0.003, **Figure 3a**).

**Figure 3:**
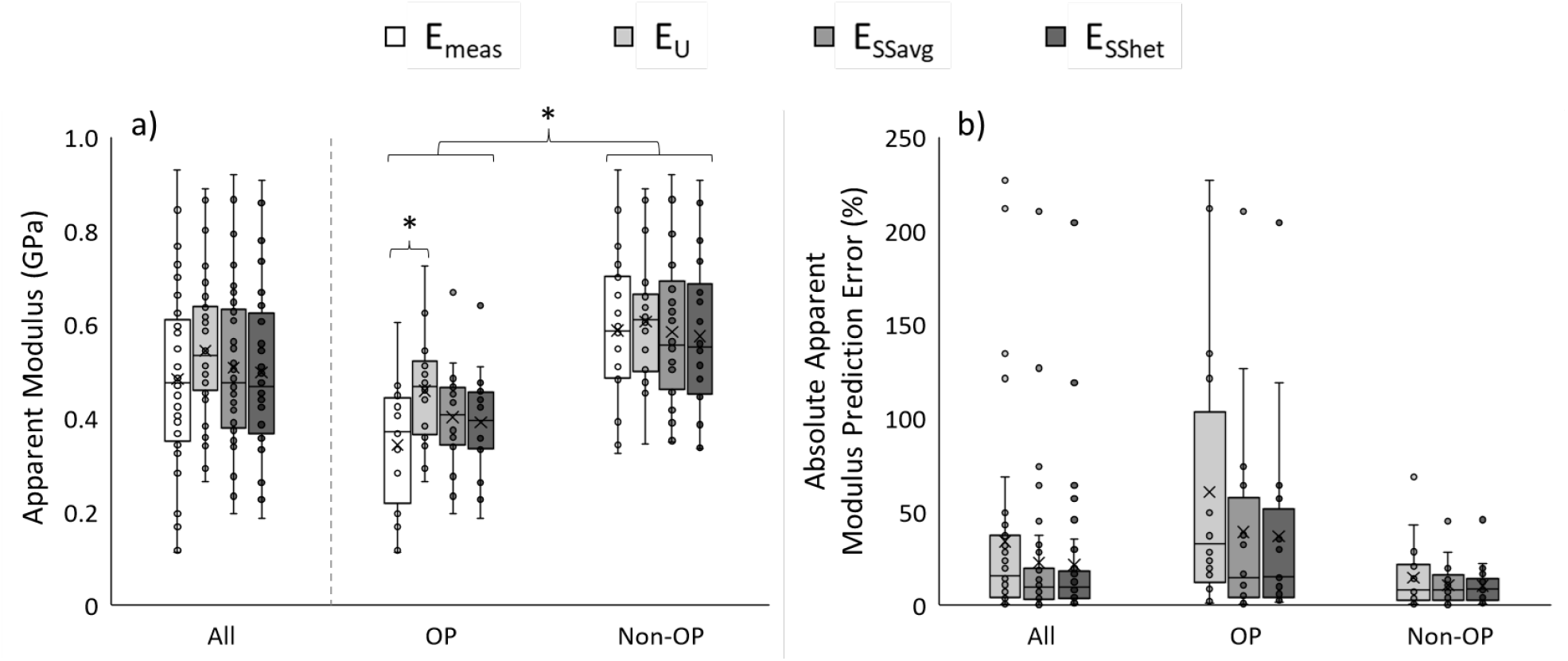
(a) Apparent modulus for each estimation method, shown for all subjects and subgroups by osteoporosis (OP) status. (b) Absolute apparent modulus prediction error, shown for all subjects and subgroups by OP status. * p < 0.05

### Apparent Modulus Prediction

Across all subjects, the absolute apparent modulus prediction error between the FE-predicted and experimentally measured values did not differ between E_U_, E_SSavg_, or E_SShet_ material distributions (38% vs. 25% vs. 24%, p = 0.21, **Figure 3b**). Absolute apparent modulus prediction error was 26% higher for osteoporotic models compared to non-osteoporotic models, but this effect was not significant after applying the Benjamini-Hochberg adjustment (p = 0.061). Additionally, the difference in modulus prediction error between osteoporotic and non-osteoporotic models did not depend on the assigned material distribution (p = 0.34). For each of the three model sets, absolute apparent modulus prediction error decreased with increasing experimentally measured modulus, appearing to conform to a negative power law relationship (**Figure 4**). Signed apparent modulus prediction error was greater for osteoporotic compared to non-osteoporotic models by 50% for E_U_ models (p = 0.034, **Figure 5**), but this difference was not significant for E_SSavg_ models (p = 0.29) or E_SShet_ models (p = 0.30). A significant bias toward overestimation of the measured modulus (signed prediction error > 0) was found only for E_U_ osteoporotic models (p = 0.0003, **Figure 5**).

**Figure 4:**
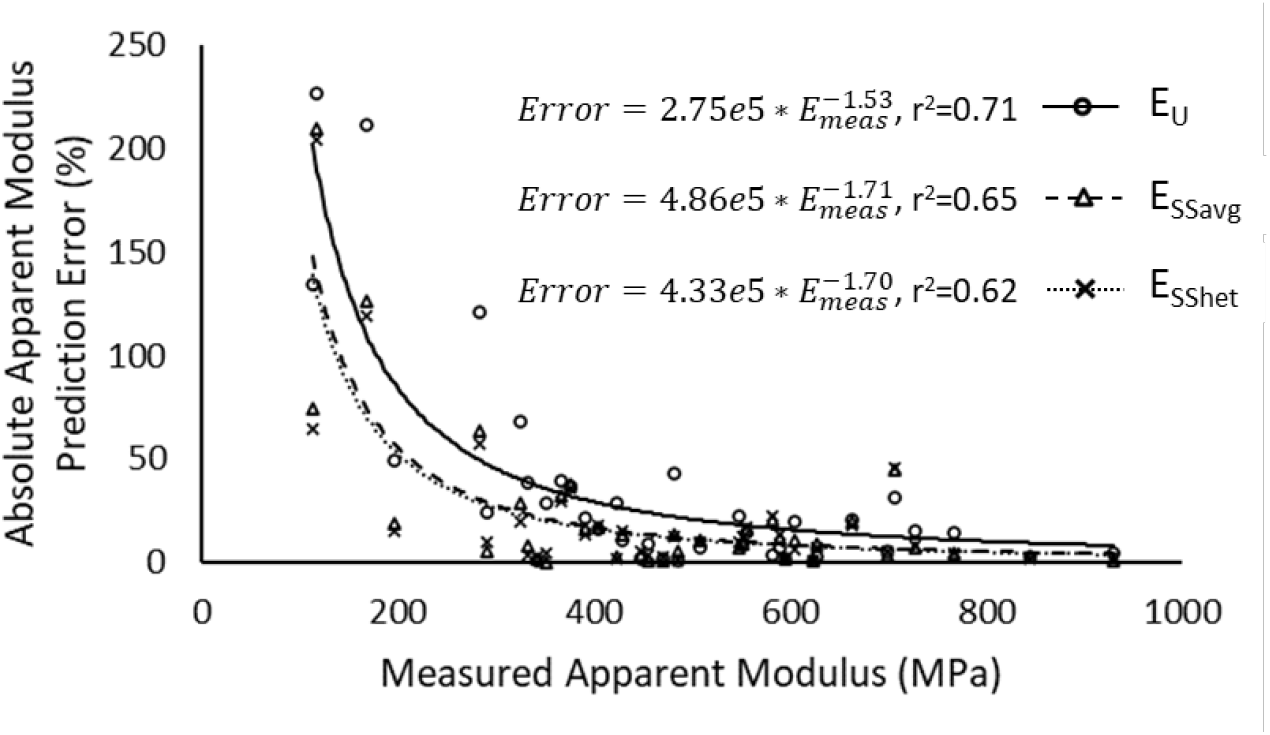
Absolute modulus prediction error for each model set, expressed as a percentage difference from the measured apparent modulus, E_meas_. Data follow a negative power law trend.

**Figure 5:**
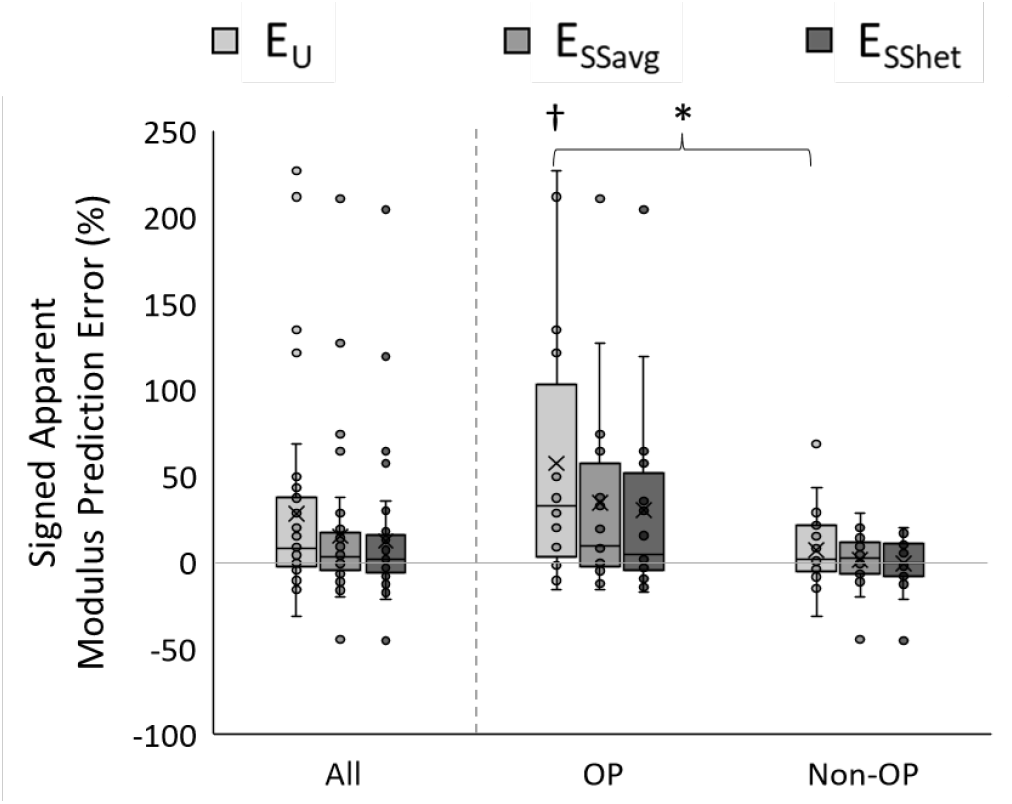
Signed apparent modulus prediction error, shown for all subjects and subgroups by OP status. * p < 0.05, † p < 0.05 vs. mean of 0.

Regardless of assigned tissue modulus distribution, FE-predicted apparent modulus was linearly related to the experimentally measured apparent modulus (p < 0.0001 for all cases) (**Figure 6a**). Overall, specimen-specific models - both E_SSavg_ and E_SShet_ models - predicted apparent modulus more reliably than E_U_ models (r^2^ = 0.61 for U_hom_, r^2^ = 0.74 for SS_hom_, and r^2^ = 0.75 for SS_het_). Among the categorical variables of sex, site, and clinical bone diagnosis, only clinical bone diagnosis had a significant effect on FE-predicted apparent modulus.

**Figure 6:**
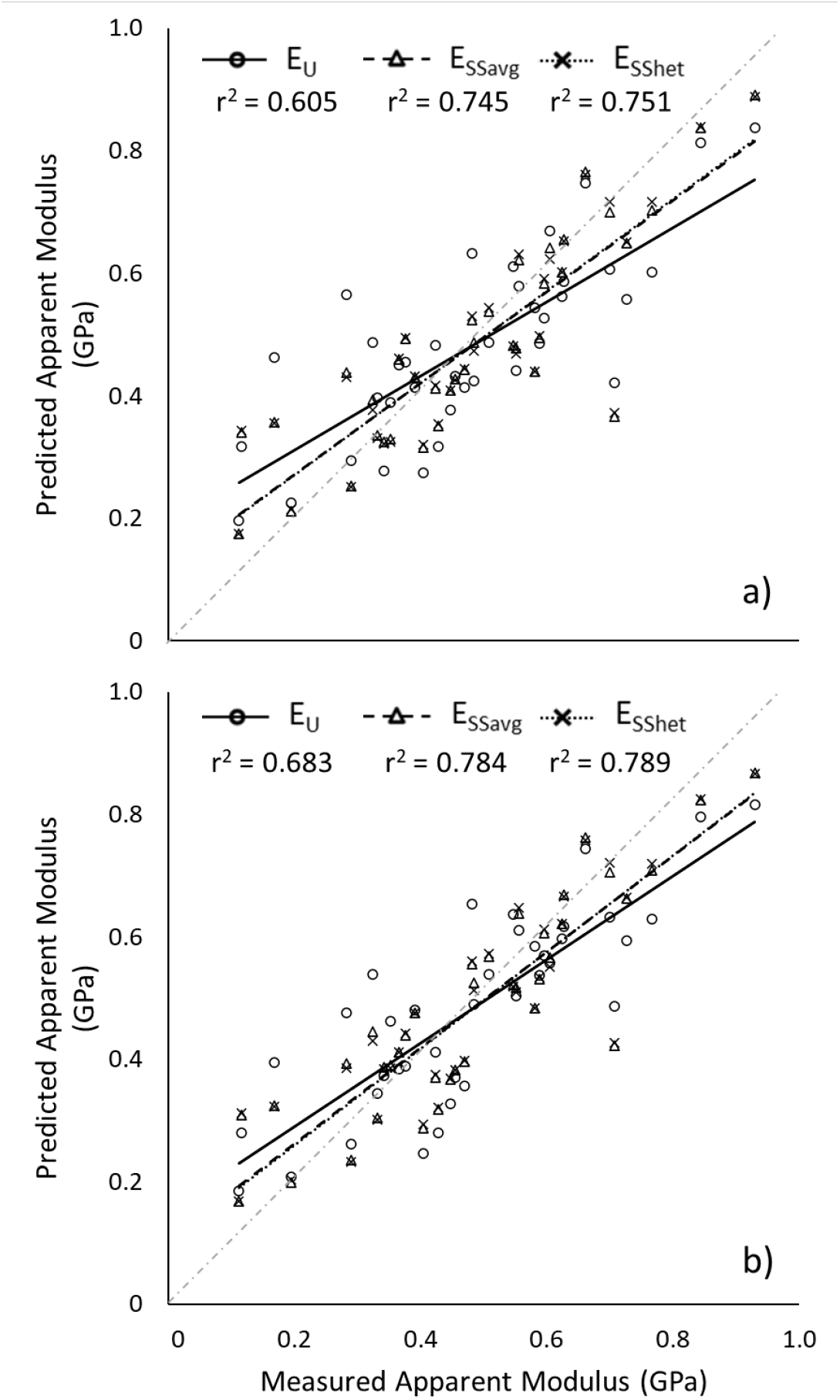
(a) FE-predicted apparent modulus for each model set was linearly related to measured apparent modulus. (b) Including clinical bone diagnosis as an explanatory variable improved r^2^ for every model set, most prominently for E_U_. Light grey lines represent unity slope.

Incorporating clinical bone diagnosis improved the prediction of experimental apparent modulus by 8% for the universal model set and 4% for both specimen-specific model sets (**Figure 6b**).

### Minimum Principal Strain

Distributions of compressive minimum principal strain (*ϵ*_*1*_) were evaluated throughout the models (**Figure 7**). Because FE analyses were linear and displacements were applied, the results for *ϵ*_*1*_ were identical in both sets of homogeneous models (i.e., E_U_ and E_SSavg_) and thus were analyzed simply as *homogeneous*. To avoid artificially high strains in elements near the applied load, a 14-mm central volume of interest was initially considered for analysis. Because the proportion of high strain elements was not different between a full-size volume and this central volume for either homogeneous (p = 0.2) or heterogeneous (p=0.9) models, all elements were included for strain analyses. For clarity, the values of compressive strain are presented in terms of the absolute value.

**Figure 7:**
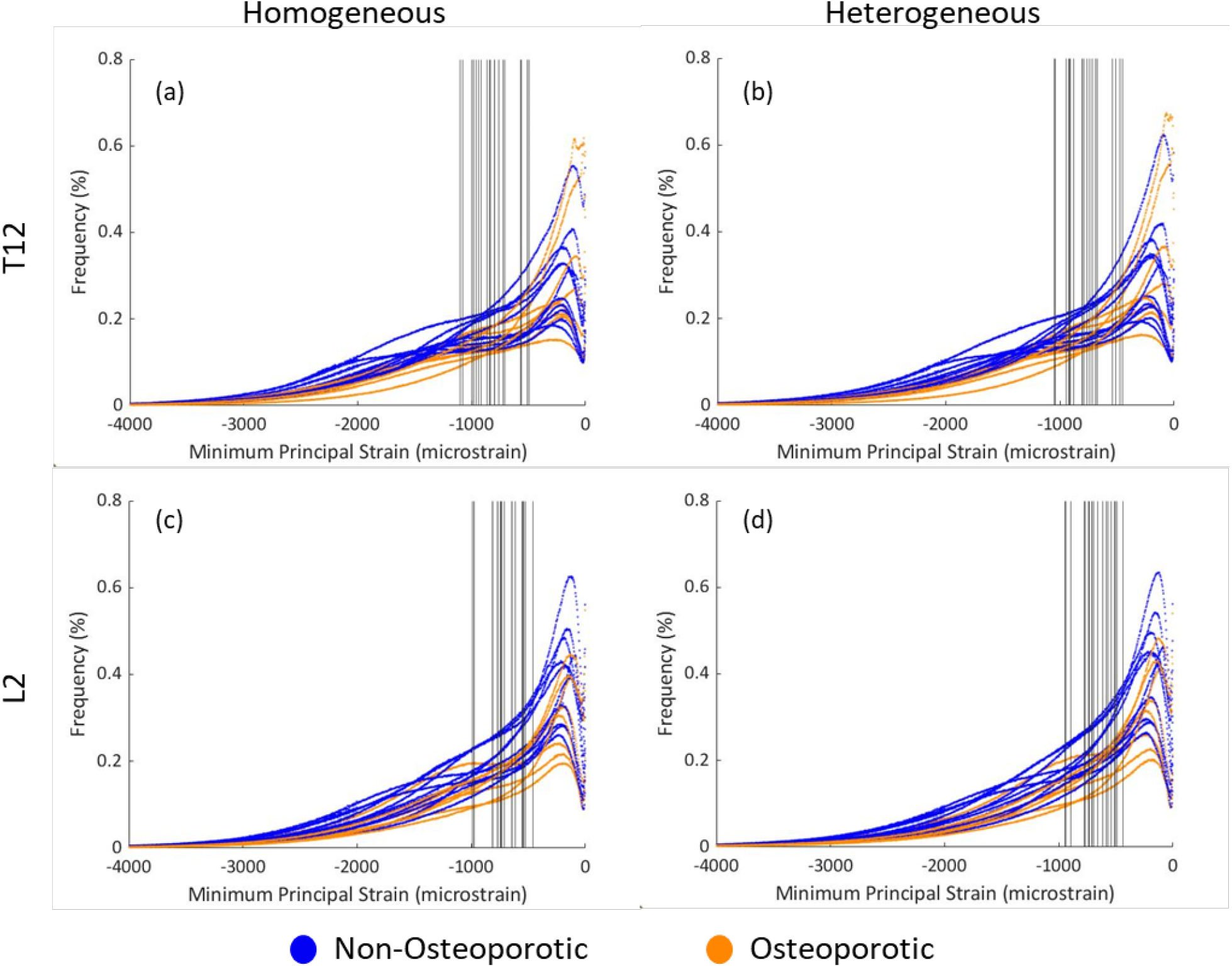
Distributions of tissue minimum principal strains (*ϵ1*) throughout the entire bone volume for homogeneous models (E_U_ or E_SSavg_) (a,c) and heterogeneous models (E_SShet_) (b,d) for all subjects, separated by T12 (a,b) and L2 bone sites (c,d). Vertical lines indicate the median value of *ϵ*_1_ for each subject.

Distributions of *ϵ*_*1*_ were similar for homogeneous and heterogeneous material models (**Figure 7**). However, element strain trended slightly lower in heterogeneous models, with median *ϵ*_*1*_ lower by 41 microstrain (p = 0.063, **Figure 8**) and 1.3% fewer high-strain elements (p = 0.086). However, when subjects were segmented by bone site, neither the T12 or L2 subset displayed any significant differences in *ϵ*_*1*_ between homogeneous and heterogeneous models. Looking at the effect of bone site, compared to T12 bone cores, L2 cores had a median *ϵ*_*1*_ that was lower by 98 microstrain (p < 0.0001, **Figure 8**), 3.8% more low-strain elements (p = 0.0003), and 3.6% fewer high-strain elements (p < 0.0001).

## 4. Discussion

This study is the first to elucidate the role of tissue modulus distribution in predicting the mechanics of osteoporotic and non-osteoporotic bone. In addition, this work offers several advances over prior FE studies investigating the effect of varying material distributions (Bourne and van der Meulen, 2004; Hammond et al., 2018; Jaasma et al., 2002; Knowles et al., 2019; Nagaraja et al., 2007; van der Linden et al., 2001; van Ruijven et al., 2007) by collectively including human samples, a larger sample size, a more clinically relevant osteoporotic fracture site, and a comparison of specimen-specific homogeneous and specimen-specific heterogeneous cases. Although the relative importance of specimen-specificity and heterogeneity in improving model performance has been in dispute (Bourne and van der Meulen, 2004; Hammond et al., 2018; Jaasma et al., 2002; Renders et al., 2011, 2008; van der Linden et al., 2001; van Ruijven et al., 2007), our results support specimen-specificity as a critical factor for evaluating the linear elastic response of human vertebral cancellous bone under uniform compression. For 24 out of 38 bone cores included in this study, the computed mean tissue modulus was lower than the assumed value of 20 GPa for the universal modulus models (**Table 1**), supporting the notion that models with specimen-specific tissue material properties may more accurately assess the structural competence of cancellous bone. Indeed, models with a specimen-specific tissue modulus substantially improved the overall prediction of apparent modulus compared to models that assumed a universal modulus within and across subjects. Notably, universal tissue modulus models tended to overestimate apparent modulus for osteoporotic subjects, while specimen-specific tissue modulus models did not. Among specimen-specific models, those that incorporated material heterogeneity performed only marginally better for predicting apparent elastic modulus than those with average material properties. Similarly, while mean TMD was strongly associated with apparent elastic modulus for these bone cores, the degree of heterogeneity indicated by TMD variability was only marginally associated.

Our study demonstrates that the simple inclusion of specimen-specific tissue material properties enabled robust model performance across subjects with varying degrees of osteoporotic bone loss and of both sexes. For osteoporotic bone, homogenization of material properties has been reported (Vennin et al., 2017) and could suggest a greater importance of average material properties compared to heterogeneity. However, our results support average material properties as the greater influence despite osteoporotic samples showing no evidence of homogenization. Alternatively, our findings could be attributed to the alignment of the applied displacement with the anatomical superior-inferior axis. For vertebral bone, trabeculae that align with this axis of compression are the main contributors to apparent modulus (Liu et al., 2006), and trabecular axial stiffness depends on average tissue density but *not* the density distribution, while bending stiffness depends on both (van der Linden et al., 2001). Therefore, the greater influence of average material properties for predicting apparent modulus is reasonable under axial compression. Models that incorporated specimen-specificity also reduced overestimation of apparent modulus by 16% for females and 12% for males, compared to models with a universally applied tissue modulus. This disparity may be explained by greater susceptibility of women to osteoporotic bone loss than men. Sex-specific changes in vertebral bone mineral content (BMC) with age may also contribute, with BMC declining in aging women but increasing in aging men (Matsui et al., 2012).

Specimen-specific models in this study were comparable to models and physical specimens from prior studies. The mean tissue elastic modulus for heterogeneous models included in this study (14.5-22.2 GPa) fell within the range of mean indentation modulus in human bone tissue examined by nanoindentation (7-32 GPa) (Donnelly et al., 2006; Hoffler et al., 2005; Remache et al., 2020; Rho et al., 1999; Rodriguez-Florez et al., 2013; Wang et al., 2006; Zysset et al., 1999). The range of predicted apparent modulus from our models (0.18-0.91 GPa) likewise fell within the range computed in previous finite element studies of human cancellous bone (0.15-2.1 GPa) (Hollister et al., 1994; Liu et al., 2013, 2008; Renders et al., 2008; van Lenthe et al., 2006; van Rietbergen et al., 1995). Furthermore, prior work involving human vertebral cancellous bone models reported reductions in apparent modulus of 2.5-5% when models with a universal tissue modulus were reassigned heterogeneous tissue moduli (Jaasma et al., 2002; van der Linden et al., 2001), similar to our results of 6% reduction for models with a specimen-specific average tissue modulus and 8% reduction for heterogeneous tissue moduli.

Though tissue heterogeneity may not strongly affect apparent modulus in human vertebral bone, it may allow models to replicate other important functions of the trabecular lattice structure, such as the ability to concentrate strain in regions with relatively low importance to load bearing. Overall, heterogeneous models in this study had a lower mean tissue modulus than models assigned a universal tissue modulus but simultaneously exhibited slightly lower strain. The idea that tissue heterogeneity influences the spatial distribution of tissue strain is further supported by a prior study in bovine tibiae, which found that incorporating heterogeneity increased the correlation strength between element strain and tissue microdamage identified through histology (Nagaraja et al., 2007). These findings suggest that tissue heterogeneity may also be an important factor influencing the behavior of sacrificial trabeculae observed by previous authors (Hayes and Carter, 1976; Liu et al., 2009; Torres et al., 2019; van der Linden et al., 2001). These trabeculae orient transversely to applied loads, contributing little to the apparent modulus but preserving structural integrity by absorbing the energy of extreme loads through plastic bending. These subtle behaviors of the cancellous bone lattice may not be essential for determining apparent modulus but are likely far more important for biomechanical phenomena that take place at the microstructural scale, such as microdamage initiation and propagation, fatigue failure, and strain-driven remodeling. Heterogeneous models consistently provide more conservative estimates of apparent modulus than models assigned a specimen-specific average modulus, explain more variability in measured apparent modulus, and are more representative of bone tissue composition and function *in vivo*; therefore, a shift toward specimen-specific heterogeneous models as the standard for FE-based bone research will most likely be beneficial.

A limitation of our bone finite element models was the use of modulus-density relationships derived from apparent-level properties rather than tissue-level measurements of true intrinsic material properties. Commonly, apparent-level tissue properties are used in the process of assigning tissue-level properties to finite elements, either through direct application of a modulus-density relationship measured at the apparent level (Austman et al., 2008; Cong et al., 2011) or optimizing the tissue modulus-density relationship based on predictions of apparent modulus (Bourne and van der Meulen, 2004). Future studies should examine whether material models based on tissue-level data (e.g., from nanoindentation or spectroscopy measures of composition) may improve the prediction of cancellous bone mechanical properties, similar to a previous study in cortical bone (Kim et al., 2012). This approach may be especially important for conditions like osteoporosis that can alter tissue-level mineralization and modulus (Kim et al., 2014).

Our study had other limitations that may have affected the influence of heterogeneity in both *ex vivo* and *in silico* analyses. For *ex vivo* testing, TMD coefficient of variation was used to represent heterogeneity, but this metric alone did not explain differences in the spatial distribution of bone mineralization between or within subjects. As previously discussed, the distribution of bone mineralization may affect both local concentrations of tissue strain (Nagaraja et al., 2007) and the bending stiffness of individual trabeculae (Kim et al., 2012; van der Linden et al., 2001). For *in silico* models, we assumed purely elastic or linear element behavior. Our physical specimens did exhibit approximately perfectly elastic behavior at the *apparent-level* up to and beyond 0.25% strain, consistent with prior measurements of human vertebral cancellous bone cores under compression (Morgan et al., 2000), but we do not know whether the *tissue-level* behavior was purely elastic throughout the bone volume. Some plastic behavior may have occurred in discrete locations of high strain, which may overlap with low-density bone tissue in bending trabeculae (Hayes and Carter, 1976; Liu et al., 2009; Torres et al., 2019; van der Linden et al., 2001). Our models also assumed an isotropic modulus for all elements, despite *in vivo* bone tissue exhibiting material anisotropy with greater yield strain and modulus in compression than in tension (Reilly and Burstein, 1975). In our models, only an average of 0.04% of elements were in tension after uniform model compression, so this limitation likely had little impact on model performance.

Finally, our findings in high-resolution FE models may not have the same significance in models generated by clinical computed tomography. The voxel sizes of clinical CT images, approximately 61 μm for peripheral sites and 150 μm for spine, are considerably larger than the voxels used in this study. At clinical CT resolution, the sampling of bone material properties and architectural boundaries are subject to partial volume effects, which tend to lower the apparent modulus (Niebur et al., 1999), especially when modeling the thinner trabeculae of osteoporotic bone. However, further research to improve the performance of FE models may compensate for the drawbacks of lower resolution, and our findings suggest that incorporating specimen-specific and heterogeneous material properties in FE models of cancellous bone may constitute a step towards this goal.

## 5. Conclusions

Comparing three sets of models with increasing material complexity, we determined that the mean specimen-specific tissue modulus of vertebral cancellous bone was more important than tissue modulus heterogeneity for governing compressive elastic behavior. Nevertheless, FE models that incorporate physiological material heterogeneity were the most accurate predictors of apparent modulus overall and were robust to differences in sex and clinical bone status. Heterogeneity may also be important for enabling certain FE models to replicate strain-concentrating adaptations of bone that protect it from damage under large apparent strain and may be more influential in nonlinear FEA aimed toward understanding the failure mechanisms of bone. Models that assumed a universal homogeneous tissue modulus were poorly suited for predicting apparent modulus, due to overestimation of osteoporotic bone stiffness. While these findings provide insight into the types of models that may prove useful for clinical bone evaluation in the future, the difference in imaging resolution between clinical CT and micro-CT remains a barrier, especially for osteoporotic bone. The utility of finite element models for elucidating factors driving cancellous bone mechanical behavior depends critically on mimicking the physical structure, material, and function of bone. Our findings may help improve the accuracy of future FE-based research, particularly related to osteoporotic bone.

## 6. Acknowledgments

We thank Dr. Kenneth Mann (SUNY Upstate Medical University) for cadaver spines and usage of lab facilities and Dr. Donald Bartel for helpful discussions. We also acknowledge Chris Wiesen and the Odum Institute for statistics consultations and acknowledge the computing resources and consultations provided by North Carolina State University High Performance Computing Services Core Facility (RRID:SCR_022168). This work was supported by the National Institutes of Health (T32 AR007281, S10 RR014801, P30 AR046121).

